# Drift-induced selection between male and female heterogamety

**DOI:** 10.1101/141929

**Authors:** Carl Veller, Pavitra Muralidhar, George W. A. Constable, Martin A. Nowak

**Author notes:** Equal contribution (Carl Veller, Pavitra Muralidhar, George W. A. Constable).

## Abstract

Evolutionary transitions between male and female heterogamety are common in both vertebrates and invertebrates. Theoretical studies of these transitions have found that, when all genotypes are equally fit, continuous paths of intermediate equilibria link the two sex chromosome systems. This observation has led to a belief that neutral evolution along these paths can drive transitions, and that arbitrarily small fitness differences among sex chromosome genotypes can determine the system to which evolution leads. Here, we study stochastic evolutionary dynamics along these equilibrium paths. We find non-neutrality, both in transitions retaining the ancestral pair of sex chromosomes and in those creating a new pair. In fact, substitution rates are strongly biased in favor of dominant sex determining chromosomes, which fix with higher probabilities than mutations of no effect. Using diffusion approximations, we show that this non-neutrality is a result of ‘drift-induced selection’ operating at every point along the equilibrium paths: stochastic jumps off the paths return, on average, with a directional bias in favor of the dominant segregating sex chromosome. Our results offer novel explanations for the ubiquity of transitions between male and female heterogamety and the preponderance of dominant major sex determining genes.

## 1 Introduction

Sex determination—the process through which an organism is directed to become either male or female—is a critical part of life history. In most animals, sex is determined genetically (Bull 1983). Among those animals with genetic sex determination, the majority exhibit *heterogametic* sex determination: the presence or absence of a sex-specific chromosome serves as the trigger for sexual differentiation (Bull 1983; Beukeboom and Perrin 2014). Depending on whether the sex-specific chromosome is in males or in females, the system is, respectively, male heterogamety (XX females, XY males) or female heterogamety (ZW females, ZZ males).

The system of heterogamety is a fundamental genetic property of a species. It is therefore surprising that it is evolutionarily very labile, with transitions between male and female heterogamety having occurred frequently in amphibians (Hillis and Green 1990), reptiles (Ezaz et al. 2009; Pokorna and Kratochvíl 2009), and fishes (Mank et al. 2006; Mank and Avise 2009; Ezaz et al. 2006), as well as in invertebrates (Kaiser and Bachtrog 2010; Vicoso and Bachtrog 2015; Becking et al. 2017). A striking example of a recent transition is found in the frog *Rana rugosa*: populations in northern Japan exhibit female heterogamety while populations in southern Japan exhibit male heterogamety (Nishioka et al. 1993; Miura et al. 1998).

In a classic theoretical study, Bull and Charnov (1977) showed that continuous paths of population genetic equilibria can be found between male and female heterogamety. States along these paths are equilibria in the sense that the evolutionary dynamics of an infinite, randomly mating population are stationary at them when all genotypes are equally fit. Intermediate states along the paths involve the presence of multiple genotypes for each sex, a situation observed in several species, including the platyfish *Xiphophorus maculatus* (Kallman 1965, 1968), the threespine stickleback *Gasterosteus aculeatus* (Kitano et al. 2009), the blue tilapia *Oreochromis aureus* (Lee et al. 2004), a Lake Malawi cichlid *Metriaclima pyrsonotus* (Ser et al. 2010), the western clawed frog *Xenopus tropicalis* (Roco et al. 2015), the housefly *Musca domestica* (Hiroyoshi 1964; Franco et al. 1982; Feldmeyer et al. 2008), and the Hessian fly *Mayetiola destructor* (Benatti et al. 2010).

Two of these equilibrium paths are of particular interest. The first, which we shall refer to as ‘model 1’, governs those transitions between male and female heterogamety that involve the same pair of sex chromosomes (Bull and Charnov 1977) (see Figures 1 and 3A). The second, which we shall refer to as ‘model 2’, governs those transitions between male and female heterogamety that involve different pairs of sex chromosomes, i.e., where the sex chromosome pair in one system is autosomal in the other, and vice versa (Scudo 1964, 1967; Bull and Charnov 1977) (see Figures 2 and 3B).

**Figure 1:**
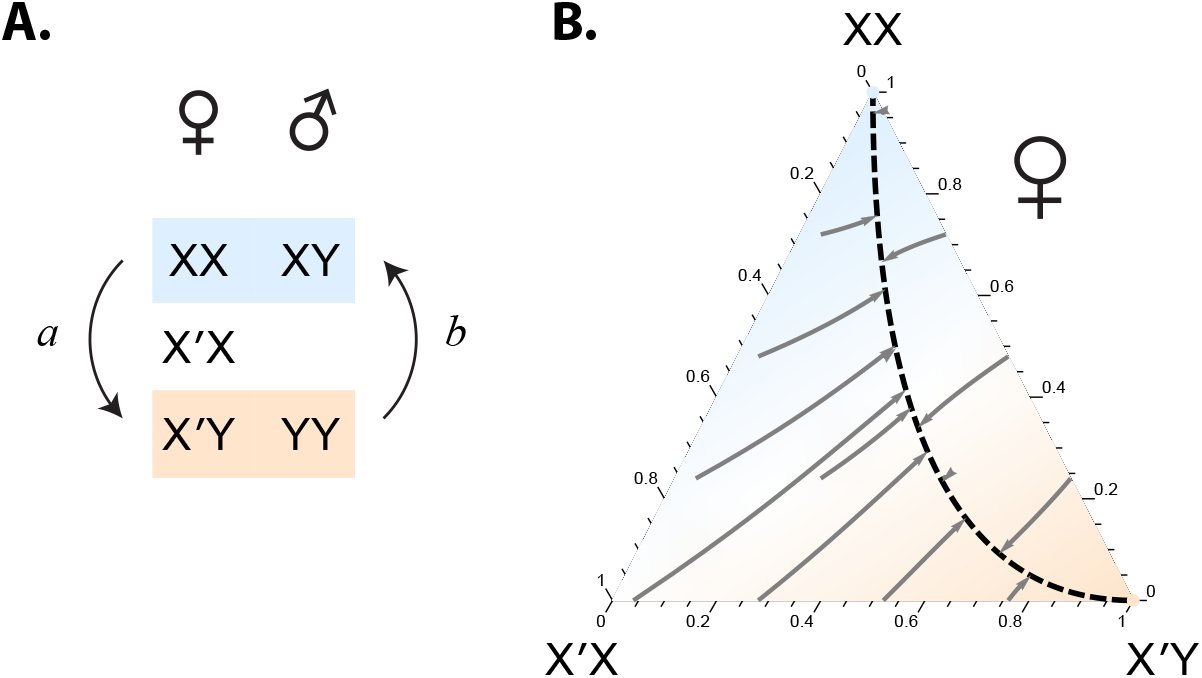
Transitions between male and female heterogamety without a change in the sex determining locus (model 1). (A) In model 1, heterogametic transitions involve intermediate systems of five sexual genotypes. In the system depicted here, there are three female and two male genotypes. Transition *a* is from male to female heterogamety, and involves fixation of the dominant feminizing X′ chromosome (mutated from an X). The reverse transition, *b*, is from female to male heterogamety, and involves fixation of the recessive feminizing X chromosome (mutated from an X′). A symmetric path, with two female and three male genotypes, also exists (see Figure 3A). (B) The equilibrium path (dashed line) governing transitions *a* and *b*, restricting attention, for ease of visualization, to the frequencies of the three female genotypes when the sex ratio is 1/2 (as it is at every point on the equilibrium path). Some deterministic trajectories to the equilibrium path are displayed with arrowed grey lines. Color shading indicates frequency of homogametic females (blue) relative to heterogametic females (orange). The equation for the equilibrium path is given in Eq. (1).

**Figure 2:**
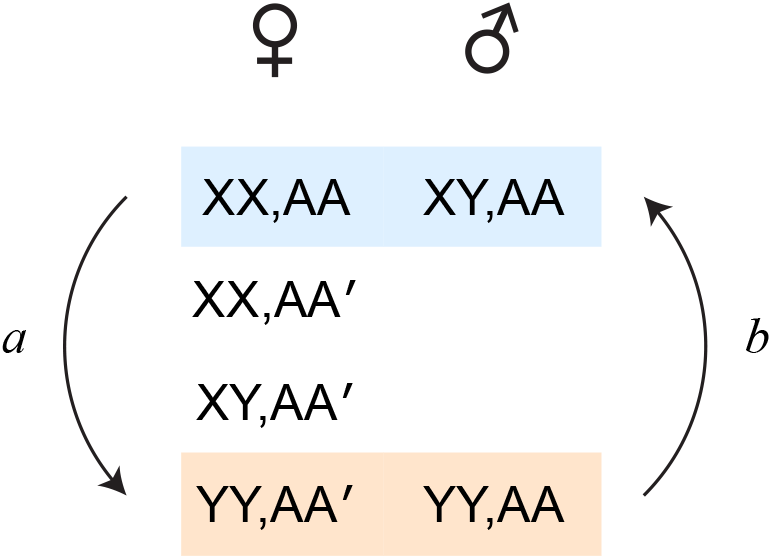
Transitions between male and female heterogamety with a change in the sex determining locus to a previous autosome (model 2). Model 2 transitional systems involve six sexual genotypes. In the system illustrated here, there are four female and two male genotypes. Transition *a* involves fixation of the dominant feminizing A′ chromosome (mutated from an A), and cause the previous sex chromosome Y to become an autosome. Transition *b* involves fixation of the recessive feminizing X (mutated from a Y), and causes the previous sex chromosome A to become an autosome. The equilibrium path of this system is given in Eq. (2). A symmetric path, with two female and four male genotypes, also exists (see Figure 3B).

**Figure 3:**
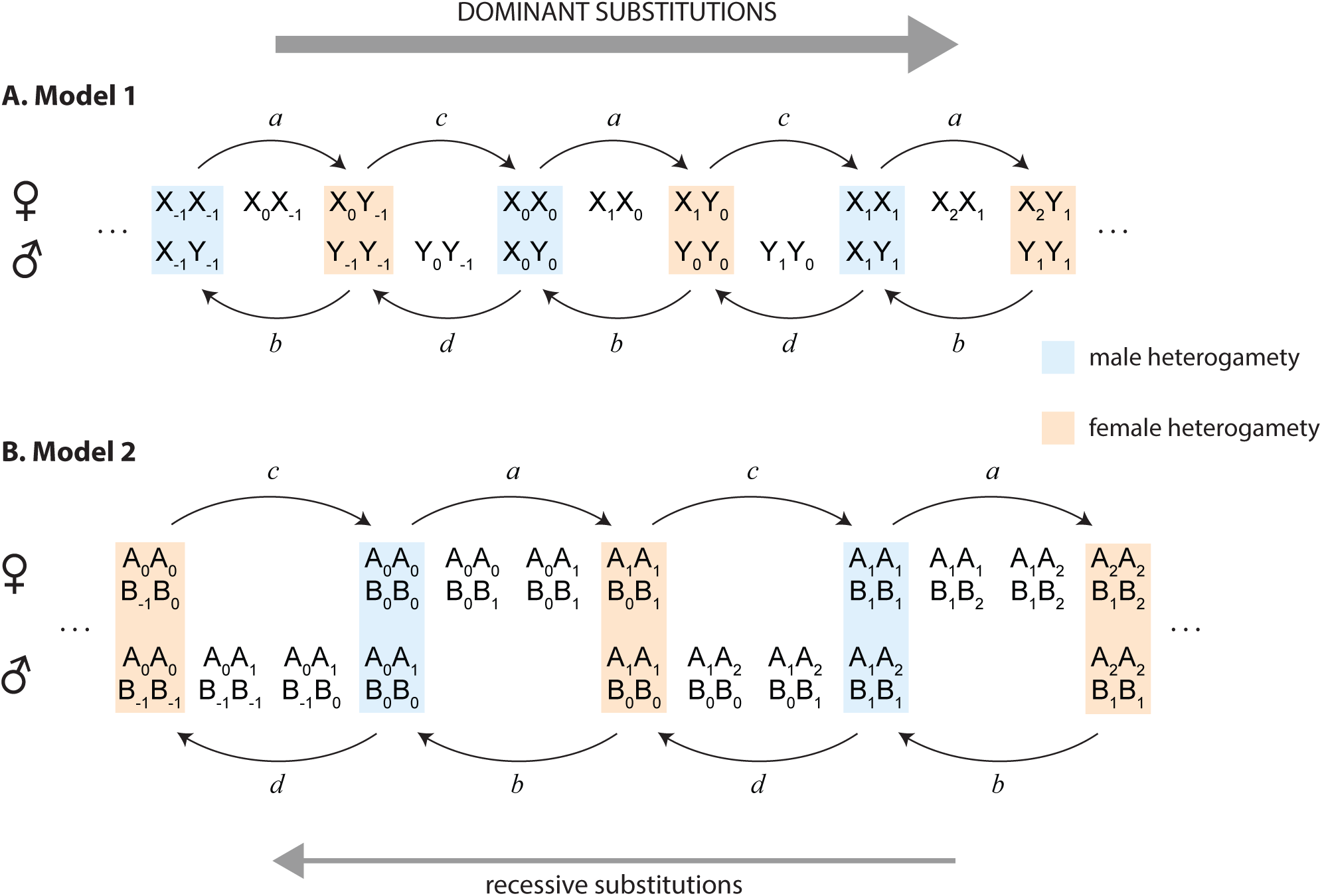
A general representation of the substitutions involved in changing the heterogametic system, both for model 1 transitions (A) and for model 2 transitions (B). In both models, numerical subscripts reveal regularities in the progression of mutations whose fixations switch the system of heterogamety. (A) For model 1, because the sex chromosome locus does not change, we have used the usual sex chromosome labels X and Y. Transitions between heterogametic systems that involve the arrival and fixation (‘substitution’) of new dominant sex determining mutations are labeled *a* and *c*. *a* is from male to female heterogamety, and involves the substitution X*_n_* → X_*n*+1_; *c* is from female to male heterogamety, and involves the substitution Y_*n*_ → Y_*n*+1_. Transitions that involve the substitution of new recessive sex determining mutations are labeled *b* and *d*. *b* is from female to male heterogamety via the substitution X_n_ → X_*n*−1_; *d* is from male to female heterogamety, via the substitution Y*_n_* → Y_*n*−1_. (B) For model 2, we have used the arbitrary letters A and B to denote the two separate chromosomes/unlinked loci: one switches to a sex chromosome, and the other switches to an autosome, each time the heterogametic system changes. Again, *a* and *c* are used to label heterogametic transitions involving substitution of new dominant sex determining mutations; *a* from male to female heterogamety via substitution of the mutation B*_n_* → B_*n*+1_ on a previous autosome, and *c* from female to male heterogamety via substitution of the mutation A*_n_* → A_*n*+1_ on a previous autosome. *b* and *d* refer to the reverse transitions, involving substitution of new recessive sex determining mutations; *b* from female to male heterogamety via substitution of the mutation A*_n_* → A_*n*−1_ on a previous autosome, and *c* from male to female heterogamety via substitution of the mutation B*_n_* → B_*n*−1_ on a previous autosome. We find in both models that, when all sexual genotypes are equally fit, dominant sex determining mutations substitute at a higher rate than recessive sex determining mutations—with reference to the present figure, we find a bias towards rightward transitions of the heterogametic system.

The existence of these deterministic equilibrium paths has led to a belief that neutral drift along them in finite populations could be responsible for transitions between male and female heterogamety (Bull 1983, 1987; van Doorn 2014), an important baseline model for such transitions (van Doorn 2014). Moreover, arbitrarily small fitness differences between the sexual genotypes can eliminate the equilibrium paths under deterministic evolutionary dynamics, and render one of the heterogametic systems stable and the other unstable (Bull and Charnov 1977). This has led to a belief that small fitness differences alone can determine which transitions are possible (Bull 1983, 1987; van Doorn 2014).

Here, we examine these claims in the context of finite-population evolutionary dynamics. Using both stochastic simulations and analytical approximations based on the removal of fast variables, we estimate the fixation probabilities of the various sex determining mutations along the two equilibrium paths, starting from a state of simple heterogamety. We find that evolution along these ‘neutral’ equilibrium paths is not neutral at all, instead showing a clear bias in favor of dominant sex determining mutations. Selection for otherwise deterministically neutral genotypes has previously been recognized in other settings (Parsons et al. 2010; Lin et al. 2012; Kogan et al. 2014; Constable et al. 2016; Newberry et al. 2016; Chotibut and Nelson 2017). Perhaps the most prominent example is Gillespie’s criterion (Gillespie 1977), which states that if the reproductive rates of two genotypes have equal arithmetic mean but different variance, then the genotype with lower variance will be selected for, owing to higher geometric mean fitness. A key difference is that, in the neutral models we study, we assume no *a priori* differences in the reproductive rates of the various genotypes; these instead emerge naturally in our analysis of the dynamics along and around the equilibrium paths.

When all genotypes are equally fit, we find in both models that the substitution rates of the dominant sex determining mutations (in directions *a* and *c* in Figure 3) are substantially higher than the substitution rates of recessive mutations (in directions *b* and *d* in Figure 3). Thus, in finite populations, stochastic evolutionary dynamics have a clear directionality along the equilibrium paths. In model 1, this directionality is amplified when natural selective forces are accounted for (viz., selection against individuals homozygous for a previously sex-specific chromosome), though the overall substitution rates in both directions are reduced. In model 2, the bias in favor of dominant sex determining mutations conferred by drift-induced selection is reduced when these selective forces are taken into account. Thus, in model 2, when selection is weak and the population is small, the substitution of dominant sex determining mutations is more likely; when selection is strong and the population is large, the substitution of recessive sex determining mutations is more likely.

The potential for drift-induced selection in transitions between sex determining systems has previously been noted by Vuilleumier et al. (2007), who use simulations to investigate the stochastic dynamics of model 1, focusing predominantly on the effects of population structure. Our work overlaps with theirs in one particular case, that of a single deme, with no viability differences between the various genotypes. In that case, they too find non-neutral fixation probabilities for new dominant and recessive sex determining mutations, but find fixation probabilities substantially above the neutral expectation for both classes of mutation. In contrast, we find in this case that dominant sex determining mutations fix with probability above the neutral expectation, but recessive mutations fix with probability lower than the neutral expectation. The fixation probabilities that they report are also orders of magnitude different from those we report, especially as population size gets large [compare, for example, their Figure 1(a) with our Figure 4A]. They also find mean conditional fixation times that are invariant, and in some cases even decrease, as population size increases; we find mean conditional fixation times that increase linearly with population size, consistent with drift-like dynamics. We have independently simulated their population model for the case that overlaps with ours, and obtain results consistent with ours, but different from those they report. The validity of our results is supported by mathematical analysis, which also allows us to explain them in analytical detail. In addition, we also consider transitions between sex determining systems in model 2, where the sex chromosome locus is changed in the course of a transition; empirically, this scenario is possibly even more common than model 1 (Ezaz et al. 2006). Our results thus suggest that biases favoring dominant sex determining mutations may be general to transitions between male and female heterogamety.

**Figure 4:**
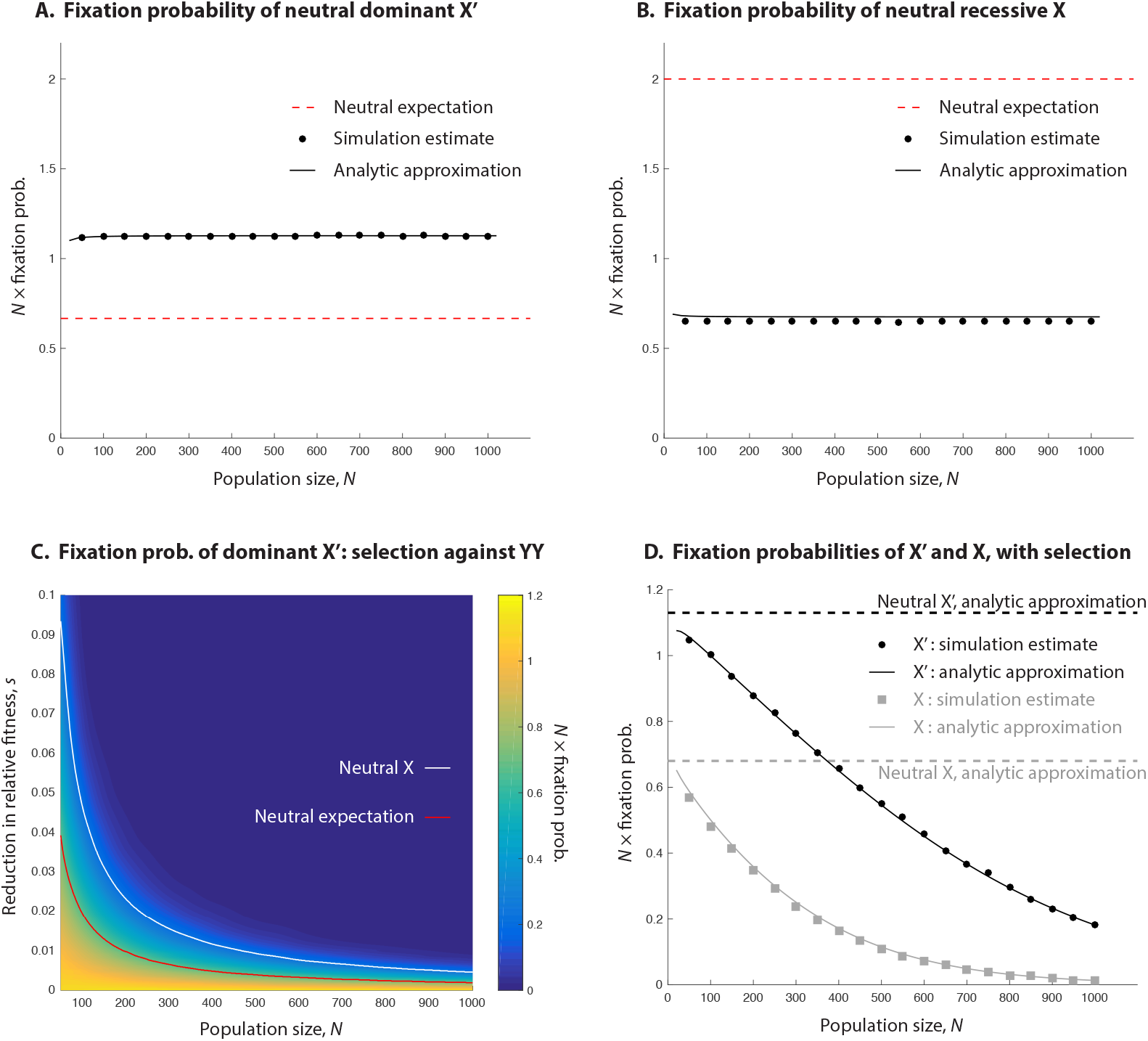
Fixation probabilities and substitution rates in Model 1 heterogametic transitions. (A, B) The fixation probability of a neutral dominant feminizing X′ arising in an XX/XY system is significantly higher than the neutral expectation (A), while the the reverse fixation probability of a neutral recessive feminizing X chromosome arising in an X′Y/YY system is much lower than the neutral expectation (B). Neutral expectations are such that the substitution rates in the two directions are equal to the mutation rate. (C) If YY individuals are selected against (fitness 1 − *s*) in the transition from XX/XY male heterogamety to X′Y/YY female heterogamety, the substitution rate of the X′ chromosome causing this transition is reduced. Still, selection needs to be sufficiently strong to reduce the substitution rate below the neutral expectation (above red line), and even stronger to reduce the substitution rate below that of the neutral X chromosome in the reverse transition (above white line). This is especially true in smaller populations. For ease of visualization, the heat map in panel C is constructed from interpolated data; the raw data are illustrated in Supplementary Information, Figure S3. Panel C is very similar to an analogous heatmap displaying our analytical results (Supplementary Information, Figure S13). (D) Fixation probabilities of the X′ and X chromosomes, in directions *a* and *b* of Figure 1A respectively, when individuals homozygous for a previously sex-specific chromosome have relative fitness reduced by *s* = 0.5% (this is in the Wright-Fisher simulations; the Moran analytical results plotted here are for a fitness reduction of 1%, which is the relevant comparison to the Wright-Fisher *s* = 0.5% owing to a two-fold increase in genetic drift under the Moran model—see Section 3.2 and Supplementary Information, Section S2.3). Though the fixation probabilities of the two chromosomes decrease when this selection operates, the relative advantage in substitution rates enjoyed by the dominant X′ over the recessive X is exacerbated.

## 2 Characterization of the equilibrium paths

### 2.1 Model 1

Consider an initial male-heterogametic system, XX/XY. Suppose now that a mutation occurs on an X chromosome that renders the feminizing tendency of the resulting chromosome—call the new chromosome X′—dominant to the masculinizing tendency of the Y (so that X′Y individuals are female). Allowing all possible male-female matings between the genotypes results in a closed system of five sexual genotypes: females can be XX, X′X, or X′Y, while males can be XY or YY (Figure 1A, system arises in direction *a*). Clearly, this system could also arise in the reverse direction: starting from a female-heterogametic system, X′Y/YY, an X′ chromosome can mutate to an X chromosome with recessive feminizing tendency (Figure 1A, direction *b*). (It should of course be noted that the usual sex chromosome labels—X, Y, Z, and W—are arbitrary, so that we could just as validly label those in a female-heterogametic system X′ and Y.)

Assuming the genotypes all to have equal fitness (i.e., that the system is neutral), enumerating them in the above order (XX, X′X, X′Y, XY, YY), and letting *p_i_* be the population frequency of genotype *i*, Bull and Charnov (1977) showed that, for any value 0 ≤ *q* ≤ 1, the population state

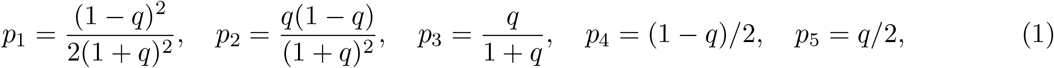

is an equilibrium in an infinite, randomly-mating population (Figure 1B). When *q* = 0, all males are XY and all females XX, so that a system of male heterogamety operates; when *q* = 1, males are YY and females X′Y, and the system is female-heterogametic. For intermediate values 0 < *q* < 1, all five genotypes are present at positive frequency.

A symmetric path exists where, from an initial female-heterogametic system ZW/ZZ, a dominant masculinizing Z′ arises from a mutated Z, and, if it reaches high enough frequency, establishes a male-heterogametic WW/WZ′ system. The reverse transition along the same path involves fixation of the recessive Z chromosome from an initial WW/WZ′ system. Intermediate states along this path involve two female and three male genotypes.

These symmetric paths, one with three female and two male genotypes (as illustrated in Figure 1), and the other with two female and three male genotypes (described in the previous paragraph), are illustrated in a general format in Figure 3A, which reveals a general numerical pattern among the mutations involved in repeated model 1 switches of the heterogametic system. In Figure 3A, among the transitions involving fixation of dominant sex determining mutations, those from male to female heterogamety are labeled *a* (as in Figure 1A), while those from female to male heterogamety are labeled *c*. Among the transitions involving fixation of recessive sex determining mutations, those from female to male heterogamety are labeled *b* (as in Figure 1A), while those from male to female heterogamety are labeled *d*. We shall use this labeling throughout for model 1 transitions.

Notice that, if there are no demographic differences between males and females, then the dynamics in directions *c* and *d* are respectively identical to those in directions *a* and *b* up to a relabeling of males and females.

For the non-neutral case, the equilibrium path connecting systems of male and female heterogamety no longer exists (Bull and Charnov 1977). The resultant dynamics in this scenario depend on the selective forces acting on each of the genotypes. Transitions in either direction involve the production of individuals homozygous for a previously sex-specific chromosome (the Y for transitions in direction *a* and the X′ and X for those in direction *b*). Sex-specific chromosomes are expected to accumulate recessive deleterious mutations in a heterogametic system (Charlesworth 1978; Charlesworth and Charlesworth 2000). We therefore expect YY genotypes to be selected against in transitions in direction *a*, and X′X and XX genotypes to be selected against in transitions in direction *b* (in direction *b*, the X chromosome is created simply by mutation at a major sex determining locus on the X′ chromosome; the X is therefore expected to carry the same deleterious mutations that the X′ does).

### 2.2 Model 2

Begin with a male-heterogametic system XX,AA/XY,AA, where A is initially autosomal. Now suppose that a mutation occurs on an A chromosome that confers on the resultant chromosome, A′, a feminizing tendency dominant to the masculinizing tendency of the Y (so that XY,AA′ and YY,AA′ individuals are female). All possible matings then yield a closed system of six sexual genotypes, females being XX,AA, XX,AA′, XY,AA′, or YY,AA′, and males being XY,AA or YY,AA (Figure 2, direction a). Again, this system could arise in the reverse direction as well, starting from the female-heterogametic system YY,AA′/YY,AA and introducing, as a mutated Y chromosome, a recessive feminizing X (Figure 2, direction *b*).

Scudo (1964, 1967) and Bull and Charnov (1977) showed that, when all genotypes are equally fit (i.e. when the system is neutral), enumerating them in the above order (XX,AA, XX,AA′, XY,AA′, YY,AA′, XY,AA, YY,AA), and writing *p_i_* for the frequency of genotype *i*, a continuous path of equilibria,

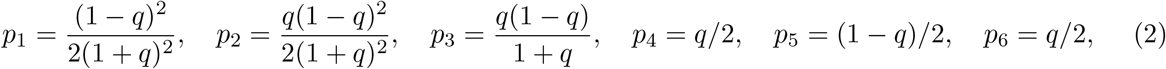

connects male heterogamety at one end (XX,AA/XY,AA; *q* = 0) with female heterogamety at the other (YY,AA′/YY,AA; *q* = 1), with intermediate equilibria (0 < *q* < 1) involving all six genotypes.

Notice that, if the former system transitions to the latter (Figure 2, direction *a*), the Y chromosome becomes autosomal, and the previous autosome A becomes a sex chromosome. Notice too that this transition involves the production of YY individuals. On the other hand, the reverse transition (Figure 2, direction *b*) does not involve the production of individuals homozygous for the previously sex-specific chromosome (here, the A′ chromosome), an important difference.

A symmetric equilibrium path exists that connects a female-heterogametic system AA,ZW/AA,ZZ with a male-heterogametic system AA′,WW/AA,WW via an intermediate system with two female and four male genotypes. Transitions from female to male heterogamety along this path involve production of WW individuals, the W having been female-specific in the original system of female heterogamety. But transitions from male to female heterogamety along this path do not involve the production of A′A′ individuals, the A′ having been male-specific under male heterogamety. That heterogametic transitions along standard equilibrium paths are possible without the production of WW or YY individuals (using the standard labeling) is an important fact often forgotten in the literature on evolutionary transitions between sex determining mechanisms.

These symmetric equilibrium paths, one with four female and two male genotypes (as in Figure 2), and the other with two female and four male genotypes (as described in the previous paragraph), are illustrated in a general way in Figure 3B. There, among model 2 transitions involving fixation of dominant sex determining mutations, those from male to female heterogamety are labeled *a*, while those from female to male heterogamety are labeled *c*. Among the reverse transitions involving fixation of recessive sex determining mutations, those from female to male heterogamety are labeled *b*, while those from male to female heterogamety are labeled *d*. Transitions *b* and *d* do not involve the production of individuals homozygous for a previously sex-specific chromosome, while transitions *a* and *c* do. We shall use this labeling throughout for model 2 transitions.

Again, notice that, if there are no demographic differences between males and females, then the dynamics in directions *c* and *d* are respectively identical to those in directions *a* and *b* up to a relabeling of males and females.

## 3 Methods

The equilibrium paths described in the previous section arise under deterministic evolutionary dynamics. Our aim is to study stochastic evolution along and around these paths. To do so, we employ Monte Carlo simulations to estimate the substitution rates of the various mutations along the paths, and approximation techniques to analytically investigate the results of these simulations.

For computational efficiency, our simulations are of a non-overlapping generations Wright-Fisher process (Wright 1931; Fisher 1930; Hartl and Clark 2007). For mathematical tractability, on the other hand, our analytical treatment considers models of the overlapping generations Moran type (Moran 1958). Agreement of the results under the two processes will demonstrate their robustness to the consideration of overlapping or non-overlapping generations.

### 3.1 Monte Carlo simulations

For both model 1 and model 2, we simulate a population of constant size *N*, which comprises males and females, and evolves according to a sexual Wright-Fisher process (Wright 1931; Fisher 1930; Hartl and Clark 2007). Each generation, males and females form mating pairs, *N* in total. An individual can be in more than one pair, and the probability that an individual is in a given pair is proportional to that individual’s fitness relative to other members of its sex (and is independent across pairs). Each mating pair produces a single offspring, whose sexual genotype (and therefore sex) is determined by randomly choosing a gamete from each of its parents. The sex chromosomes of heterogametic individuals are assumed to segregate in a Mendelian fashion. After offspring production, all individuals in the parental generation die. (This model is equivalent to one in which each male contributes a large number of sperm, proportional to his fitness, to a common sperm pool, each female contributes a large number of eggs, proportional to her fitness, to a common egg pool, and each of *N* offspring is then formed by drawing a random sperm and a random egg from the respective pools.)

As a baseline for both models, we consider the case where each genotype is equally fit (the ‘neutral’ case), i.e., where each individual within a sex is equally likely to be chosen to be in a given mating pair. Thereafter, we focus on cases where individuals that are homozygous for a previously sex-specific chromosome suffer a selective disadvantage: each such individual is only 1 − *s* as likely as other members of its sex to be chosen to be in a given mating pair.

In all cases, we assume no population structure (mating is random), and no demographic differences between males and females in our simulations. Relaxing these assumptions is an important direction of future research, and would be aided by parallel theoretical developments in the general study of drift-induced selection.

### 3.2 Diffusion approximations

In the Moran formulation, we likewise consider a discrete population of *N* individuals. Males and females of each genotype (five genotypes for model 1 and six genotypes for model 2) encounter each other with a probability per unit time proportional to their frequency in the population. On encountering each other, a pair produces a single offspring, which inherits its sexual genotype from its parents in a Mendelian fashion. In order to keep the population size constant at *N*, another individual is chosen to die. In the neutral case, each individual in the population is equally likely to be chosen to die. In the case with selection, the probability that an individual is chosen to die is weighted by a normalized death probability, the inverse of that genotype’s fitness. (Notice that this form of selection, involving increased probabilities of death, is slightly different to that in our simulations, which involve reduced probabilities of being in a mating pair, or, equivalently, a reduced contribution to a common gamete pool. If there is selection on females and males simultaneously, this will lead to a quantitative difference between the models; in the Wright-Fisher simulations, pairs of males and females that are both selected against produce offspring at a reduced rate proportional to (1 − *s*)^2^, while in our Moran model these males and females die at an increased rate proportional to (1 + *s*). The difference between the resulting dynamics will therefore be of order *s*^2^ and so, as we shall see, not significant in the weak selection limit in which we shall be working.) The full set of probability transition rates, *T*(***P***|***P′***), which gives the probability per unit time of transitioning from state ***P′*** to ***P*** (where ***P*** and P’ are vectors of the number of each genotype in the population), are given for each model in Supplementary Information, Section S2.4.

The system’s dynamics can now be entirely described by a master equation (van Kampen 2007), a large set of partial difference equations for the time evolution of the probability Φ(***P***, *t*) of being in each state ***P*** at time *t*. Since these equations are difficult to analyze, we make use of a classic tool of population genetics, the diffusion approximation (Crow and Kimura 1970). Assuming *N* to be large but finite, we can transform into approximately continuous variables ***p*** = ***P***/*N* and Taylor expand the master equation to arrive at the approximate description:

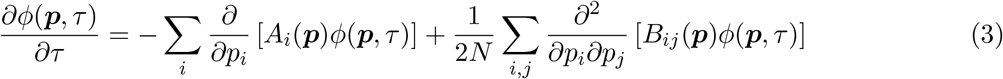

(Gardiner 2009; McKane et al. 2014), where *τ* = *t*/*N*. Here *ϕ*(***p***, *t*) is a continuous approximation of Φ(***P***, *t*) and the forms of the vector ***A***(***p***) and the matrix *B*(***p***) can be directly calculated for both models 1 and 2 from their respective transition rates (see Supplementary Information, Section S2.1). The term ***A***(***p***) primarily controls the time-evolution of the mean of *ϕ*(***p***, *t*), and can thus be interpreted as a deterministic selective term. Indeed, in the deterministic limit (*N* → ∞), the dynamics of the system are captured by the ordinary differential equations ***ṗ*** = ***A***(***p***). Meanwhile, the term *B*(***p***) controls the diffusion of *ϕ*(***p***, *t*), thus capturing the effect of genetic drift.

In comparing this diffusion equation with one for the analogous Wright-Fisher process, we must remember to take account of two scalings. Firstly, genetic drift is typically a factor 2 larger in the Moran formulation than in the Wright-Fisher formulation (Felsenstein 1971). This leads to an additional prefactor of 1/2 appearing before the *B*(***P***) in the Wright-Fisher formulation of Eq. (3). Second, reproduction occurs 4*N* times faster in the Wright-Fisher model. This is because individual reproduction events in the Moran model with sexual reproduction occur at an average rate proportional to the product of the frequency of males and females in the population (which is 1/4 at equilibrium sex ratios) whereas in the Wright-Fisher model *N* reproduction events occur per time-step. Since we have already accounted for a factor *N* in timescale in obtaining Eq. (3) (*τ* = *t*/*N*), the Wright-Fisher formulation of Eq. (3) contains an additional factor of 4 preceding all terms on the right-hand side of Eq. (3).

We wish to determine the probabilities of transitions from male to female heterogamety and vice-versa. The calculation of these quantities is not straightforward; the diffusion equation (3) governing the dynamics is non-linear in either four or five variables (for models 1 and 2, respectively). However, analytic progress can be made by utilizing results on fast variable elimination in stochastic systems (Parsons and Rogers 2015). The key to progress is in noting the following. In the deterministic neutral systems, a trajectory starting from any set of initial conditions will quickly collapse to a point on the equilibrium path of the system, Eq. (1) for model 1 (Figure 1B) and Eq. (2) for model 2, where it will then stay. In the current notation, this line is the set of solutions ***p*** to the equation ***A***(***p***) = 0. When genetic drift and selection are accounted for, the system will no longer reach and subsequently remain at a position on this line. However, if selection is weak and *N* is large (such that the rate of genetic drift, 1/*N*, is small), then the system will quickly collapse to a subspace in the vicinity of this line; it will then slowly move along this ‘slow subspace’ until the system fixes in a state of either male or female heterogamety. We exploit this separation of timescales by removing the fast transient dynamics to obtain an approximation for the system dynamics in the slow subspace. Since this approximate description in the slow subspace is in terms of a single variable, *q* [see Eqs. (1) and (2)], fixation probabilities are then straightforward to calculate.

The full calculation leading to the approximate description is provided in the Supplementary Information, Section S2.2 [also see Parsons and Rogers (2015)]. Here we simply state the key results. On removing the fast-variables, the dynamics of Eq. (3) can be approximated by

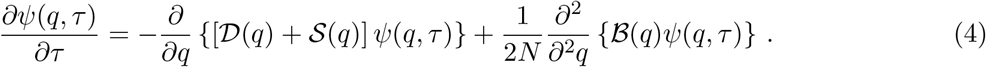

Here *ψ*(*q*, *t*) is the probability density function for *q* along the slow subspace. In a similar fashion to Eq. (3), the terms 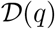 and 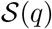 control the time-evolution of the mean of *ψ*(*q*, *t*) and can thus be interpreted as selective terms for *q* along the slow subspace. Likewise, the term 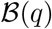 controls the diffusion of *ϕ*(***p***, *t*), and can thus be interpreted as capturing the effect of genetic drift along the slow subspace.

The equation for the slow subspace itself can be approximated by the equation for the line of equilibria, since they lie close to each other in the weak selection limit, and coincide when selection is absent. A similar approach is taken in deriving the dynamics of diploid population genetic models with selection, where it is assumed that if selection is weak, allele frequencies are approximately in Hardy-Weinberg equilibrium (Crow and Kimura 1970).

The equations for 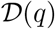, 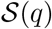, and 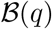 can be determined from ***A***(***p***) and *B*(***p***) along with the equation for the slow subspace. The terms 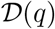 and 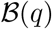 are simply the components of ***A***(*p*) and *B*(***p***) along the slow subspace (i.e., respectively, the components of deterministic selection and genetic drift in the subspace). More interesting is the term 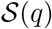, which is a selective term induced by genetic drift. Its origin can be interpreted in various ways. First, it can be graphically understood as resulting from a bias in how fluctuations taking the system off the slow subspace return to the slow subspace (i.e., fluctuations do not return, on average, to the point from which they originated on the slow subspace) (Parsons and Rogers 2015; Constable et al. 2016). Second, it can be mathematically understood as the result of making a non-linear change of stochastic variables (Risken 1989) into the system’s slow variable. Finally, it can be understood as the result of a selective pressure favoring genotypes with a lower variance in their reproductive output—Gillespie’s Criterion (Gillespie 1974; Parsons et al. 2010; Hansen 2017). In Eq. (4), however, differences in the variance of the reproductive output of the genotypes do not arise from any *a priori* assumptions about their reproductive rates (as in Gillespie’s Criterion), but instead emerge naturally from the confinement of the system to the slow subspace.

## 4 Results

### 4.1 Model 1: Transitions using the same chromosomes

In this section, we study transitions between male and female heterogamety where the sex chromosome locus is the same under both systems (Figures 1 and 3A).

#### Monte Carlo simulations

We begin with a male-heterogametic system, XX/XY, in a population of constant size N, initially with *N*/2 females (all XX) and *N*/2 males (all XY). We consider a mutation to one of the X chromosomes, rendering it an X′ chromosome, X′Y and X′X bearers of which are female. YY individuals are male. If the X′ chromosome ‘fixes’ in the population, a female-heterogametic system, X′Y/YY, is established (direction *a* in Figures 1A and 3A).

Influenced by whether the original X′ mutation occurs in oogenesis or spermatogenesis, the X′ could initially find itself in an X′X or an X′Y female. We consider both cases in our simulations, and in both begin with a population that is *N*/2 females (one of which carries an X′ chromosome) and *N*/2 males.

Initially assuming all five sexual genotypes to be equally fit, what is a reasonable null expectation for the fixation probability of this X′ chromosome? To answer this, we consider an alternative mutation on an X chromosome, which has no effect on either the fitness or the sex of its bearer. The probability that this truly neutral mutated X chromosome fixes in the ‘population’ of X chromosomes is simply the inverse of the initial census count of the X chromosome, i.e., 1/(3*N*/2). This we take to be the neutral expectation for 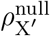, the fixation probability of the X′ chromosome, so that 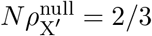.

Instead, we find in our simulations that the fixation probability of the X′ chromosome is substantially higher than this neutral expectation, with *N*_*ρ*X′_ ≈ 1.12 (Figure 4A). This is irrespective of the background on which the initial X′ chromosome finds itself (Supplementary Information, Figure S1A). The average conditional fixation time of the X′ chromosome is close to 2.4*N* for each *N* considered in our simulations, and again, this is irrespective of the mutation’s initial background (Supplementary Information, Figure S1B). A fixation time that scales linearly with *N* is strongly suggestive of drift-like evolutionary dynamics.

We now consider evolution in the other direction along the path (direction *b* in Figures 1A and 3A). We begin with an established X′Y/YY female-heterogametic system, and consider a mutation to one of the X′ chromosome that renders it an X. If this X chromosome subsequently fixes in the population of X and X′ chromosomes, an XX/XY male-heterogametic system would be established.

Here, the X mutation must occur in oogenesis, since it derives from an X′ chromosome which must have been borne by a female. Therefore, the first individual to carry the new X chromosome must be an XY male. For consistency, we begin our simulations with *N*/2 females and *N*/2 males (one of which carries the X chromosome).

Similar to before, our neutral expectation for the fixation probability of the X is just the inverse of the number of X′ chromosomes initially in the population, i.e., 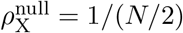, so that 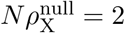. Instead, in our simulations we find the fixation probability of the X chromosome to be much lower than this neutral expectation, with *ρ*_X_ ≈ 0.65 (Figure 4B). The average conditional fixation time of the X chromosome is, like that of the X′, close to 2.4*N* for each *N* considered (Supplementary Information, Figure S2B).

Thus, the fixation probability of the X′, starting from an initial XX/XY system, is almost twice as great as the fixation probability of the X, starting from an initial X′Y/YY system. However, to properly determine whether evolution along the equilibrium path is biased in favor of the X′ chromosome, we must consider the *substitution rates* of the two chromosomes. In doing so, we assume symmetric mutation rates in the two directions, i.e., that the probability of an X chromosome mutating to an X′ is the same as the probability of an X′ chromosome mutating to an X. We also assume this mutation rate to be sufficiently small that at most one mutation segregates in the population at any given time [a common assumption (McCandlish and Stoltzfus 2014)].

Suppose this mutation rate to be *u* per chromosome per generation. Then an XX/XY population with even sex ratio produces X′ mutations at a rate of *μ*_X′_ = (3*N*/2)*u* per generation, while an X′Y/YY population produces X mutations at a third of this rate, *μ*_x_ = (*N*/2)*u* per generation. Notice that, had both the X′ and X chromosomes exhibited fixation probabilities equal to their null expectations, 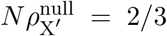 and 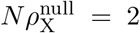, then their substitution rates would be equal, since 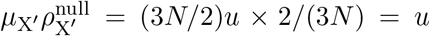 and 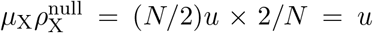. Instead, the substitution rate from an XX/XY to an X′Y/YY system (in direction *a* of Figures 1A, 3A), *μ*_X′_*ρ*_X′_, is about (3*N*/2)*u* × 1.12/*N* = 1.68*u*, while the substitution rate from an X′Y/YY to an XX/XY system (direction *b* of Figures 1A, 3A), μ_X_p_X_, is about (*N*/2)*u* × 0.65/*N* = 0.325*u*.

Thus, not only is the fixation probability of the X′ greater than its neutral expectation, and the fixation probability of the X lower than its neutral expectation, but mutational effects in fact exacerbate this asymmetry, with the substitution rate of the X′ more than *five times higher* than that of the X.

Since, in the population model we have simulated, there are no demographic differences between males and females, the transitions symmetric to those above (i.e., in directions *c* and *d* in Figure 3) occur with fixation probabilities and substitution rates equivalent to those we have estimated above (i.e., with reference to Figure 3, the fixation probabilities in directions *a* and *b* match those in directions *c* and *d* respectively).

Therefore, the most likely trajectory that the neutral dynamical system will follow in the long term is the recurrent invasion and fixation of successive *dominant* sex determining mutations, flipping the system repeatedly between male and female heterogamety. This corresponds to a bias in favor of rightward transitions in Figure 3.

We now consider the possibility that some genotypes are fitter than others. In particular, we study the case where individuals homozygous for a previously sex-specific chromosome (including mutated versions of it) are of lower fitness.

The transition from a male-heterogametic XX/XY system to a female-heterogametic X′Y/YY system, via fixation of the dominant X′ chromosome and displacement of the X, involves the production of YY males. Since the Y was sex-specific in the original XX/XY system, it may have accumulated recessive deleterious mutations that render the YY males of reduced fitness. We expect this to decrease the fixation probability of the X′ chromosome, and therefore to reduce the substitution rate in direction *a* of Figures 1A, 3A.

Assume then that YY males have relative fitness 1 — *s*, while all other sexual genotypes are of equal fitness 1. Figure 4C gives the fixation probability of the X′ chromosome (multiplied by N) for various values of *s* and *N*. As expected, the fixation probability decreases as selection against the YY genotype increases. Nonetheless, it is clear that the YY genotype can suffer appreciable fitness reductions with the X′ chromosome still fixing with probability higher than the neutral expectation. This is especially true in small populations, in which selection acts less efficiently (Lanfear et al. 2014); we would expect this to carry over to structured populations of larger size, e.g., those that are subdivided into many small demes in which drift can be an important force (Laporte and Charlesworth 2002; Whitlock 2003).

The reverse transition, from the X′Y/YY to the XX/XY system, involves the production of X′X and XX females. Since the X′ chromosome is sex-specific in the original X′Y/YY system, it may have accumulated recessive deleterious mutations, and because the X is just an X′ mutated at the sex determining locus, it too should carry these other, deleterious mutations. Therefore, both the X′X and XX genotypes should suffer reduced fitness. The substitution rate of the X, in direction *b* of Figure 1A, was very low even when no genotypes were of reduced fitness; selection against the X′X and XX genotypes severely exacerbates this disadvantage (Figures 4D and S4). Again, given the symmetry between transitions in directions *a* and *c* in Figure 3, and between transitions in directions *b* and *d*, the above results imply a bias towards *c* transitions over *d* transitions, with this bias exacerbated by selection. That is, selection generally exacerbates the bias in favor of rightward (dominant substitution) transitions in Figure 3.

#### Analytical results

To better understand these simulation results, we now study the system analytically in the diffusion limit. We first consider the neutral case. Applying the fast-variable elimination described in *Methods* and detailed in full in the Supplementary Information, Section S2, we find for the model 1 system described in Eq. (1) that the system dynamics can be approximated by Eq. (4) with 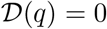 and

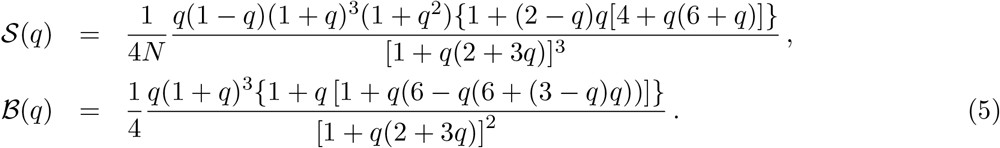

(calculations in Supplementary Information, Section S2.2.1). As we expect in this neutral scenario, there are no deterministic contributions to selection along the equilibrium path 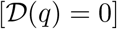. However, there is a drift-induced selection term 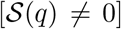. The strength of the drift-induced selection is of order 1/*N*, generated as it is by demographic fluctuations. Recalling that *q* = 0 corresponds to male heterogamety [XX/XY; *p*_1_ = 1/2, *p*_4_ = 1/2 in Eq. (1)], while *q* = 1 corresponds to female heterogamety [X′Y/YY; *p*_3_ = 1/2, *p*_5_ = 1/2 in Eq. (1)], we find that the drift-induced selection selects for the fixation of female heterogamety *at every point* on the equilibrium path [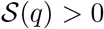 for all *q* ∈ (0,1)].

How does this drift-induced selection emerge? Essentially, in this ‘neutral’ stochastic system, demographic fluctuations in the number of genotypes constantly move the system away from the equilibrium path (to which the deterministic system is constrained). Once here, there is a selective pressure for the system to return to the equilibrium path. However, the nonlinear trajectories along which these fluctuations return, combined with the curvature of the equilibrium path, give rise to a bias in the average position to which fluctuations return. In other words, fluctuations arising at a point *q* return on average to a point *q* + *δ*(*q*) on the equilibrium path.

Mathematically quantifying this bias requires taking account of the probability distribution of fluctuations in each genotype, the form of trajectories back to the equilibrium path, and the curvature of the equilibrium path itself, each of which varies as *q* is varied. However, further intuition can be gained by decomposing 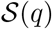 into two components: 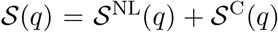. The first term, 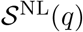, quantifies contributions to 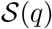 arising from the non-linearity of trajectories that take the system back to the equilibrium path. The second term, 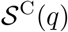, quantifies contributions to 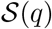 arising from the curvature of the equilibrium path itself. Plotting these terms together in Figure 5A, we see that it is the curvature of the equilibrium path that contributes most to the observed drift-induced selection.

**Figure 5:**
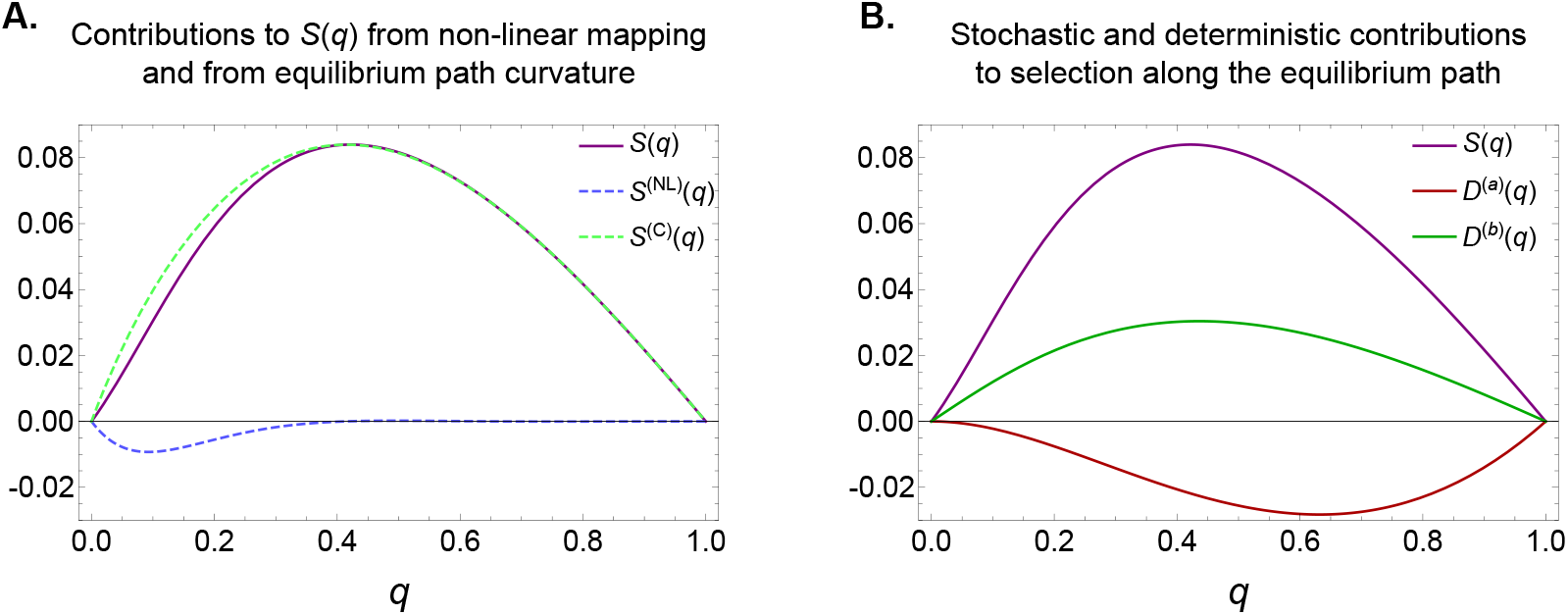
(A) In model 1, random demographic fluctuations induce a selective gradient 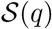 along the equilibrium path in favor of the dominant sex determining chromosome, i.e., causing *q* to increase on average [see Eqs. (4) and (5)]. 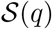 can be divided into two components. The first, 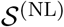, is from non-linear components of the mapping of fluctuations back to the equilibrium path; the second, 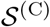, is from the curvature of the equilibrium path. In this case, it can be seen that the primary contribution to the drift-induced bias 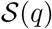 is from curvature of the equilibrium path. (B) Contributions to average dynamics along the equilibrium path arising from drift-induced selection, 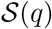, and deterministic selection against individuals homozygous for a previously sex-specific chromosome, 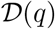 [see Eq. (4)]. Note that the form of 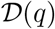 is expected to change depending on whether the initial population exhibits XX/XY male heterogamety 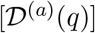 or X′Y/YY female heterogamety 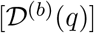 [Eqs. (7) and (8)], i.e., whether the transition is in direction *a* or direction *b* of Figure 1A. In both plots the parameters have been scaled such that *N* = *δ* = 1 in order to facilitate the comparison of contributions to selection along the slow subspace.

Standard methods can be used to numerically calculate the fixation probability of either male or female heterogamety for any initial condition *q*_0_ on the slow subspace [see Gardiner (2009); Risken (1989); and Supplementary Information, Section S2.3]. Our final task is to calculate how the initial conditions described in the previous section, those of single mutants invading a resident population, map onto initial conditions on the equilibrium path. That is, for a given ***p***_0_, we wish to determine *q*_0_. For transitions in direction *a* of Figures 1A and 3A, involving fixation of the X′ chromosome, let the initial conditions be denoted 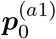 when the first X′ appears in an X′X genotype, and 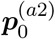 when it appears in an X′Y genotype. For transitions in direction *b*, involving fixation of the X chromosome, we denote by 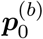 the initial condition where a single X chromosome is present in the population (necessarily in an XY male, as explained earlier). Then the respective initial conditions on the equilibrium path can be shown to be (see Supplementary Information, Section S2.3.1)

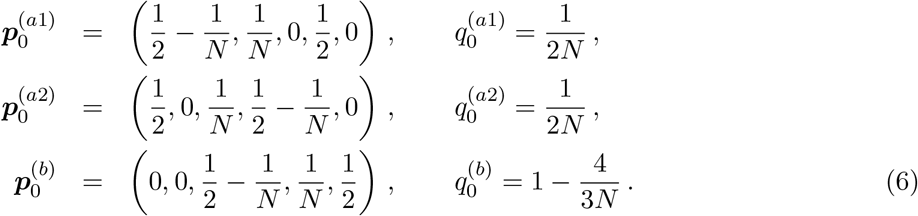

The neutral fixation probabilities can now be calculated.

The results for the fixation probabilities of the mutants are given in Figure 4A, in which we see excellent agreement between theory and simulations. In particular we find that the fixation probability of the X′ chromosome under the diffusion approximation is *N*_*ρ*X′_ ≈ 1.13 (Figure 4A). This is irrespective of the background on which the initial X′ chromosome finds itself, as both of these initial conditions lead to the same initial condition on the equilibrium path [*q*_0_ = 1/(2*N*); see Eq. (6)]. We may also compute the mean conditional fixation time of the X, which agrees well with our simulation estimates when the differences in variance between the Wright-Fisher and Moran processes are accounted for (Supplementary Information, Figure S1B). Meanwhile the fixation probability of an X chromosome is *ρ*_X_ ≈ 0.68, again in close agreement with our simulation results (Figure 4B), as is the computed mean conditional fixation time (Supplementary Information, Figure S2B).

We now consider the possibility that some genotypes are fitter than others. For transition direction *a*, assume that YY males (with frequency *p*_5_) have relative death rate Δ_5_ = 1 + *s*, while all other sexual genotypes have death rate 1. When s is small, we can still utilize fast-variable elimination to arrive at the approximate description of the system dynamics given by Eq. (4). The functions 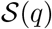 and 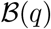 retain the forms given in Eq. (5), but now 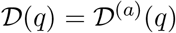, where

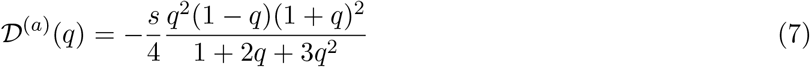

and the superscript ‘(*a*)’ denotes that this is 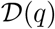 evaluated for transitions in direction *a* (Figure 5B; calculations in Supplementary Information, Section S2.2.1). This term is of order s as this is the deterministic contribution to the dynamics along the slow subspace.

For the reverse transition *b*, assume that XX and X′X females (with frequencies p_1_ and p_2_) have relative death rates Δ_1_ = 1 + *s* and Δ_2_ = 1 + *s* respectively, while all other sexual genotypes have death rate 1. Again utilizing fast variable elimination, we find 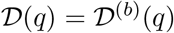, where

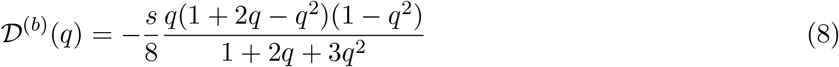

and the superscript ‘(*b*)’ denotes that this is 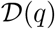 evaluated for transitions in direction *b* (Figure 5B; calculations in Supplementary Information, Section S2.2.1).

We are now in a position to calculate the respective fixation probabilities of mutations arising in directions *a* and *b*. We find that, while the fixation probability of X′ in direction *a* is of course lower than in the neutral case (since YY is selected against), the relative reduction of the fixation probability of X in direction *b* is even higher (Figure 4D). Intuitively, this can be understood as a result of the fact that only a single genotype is selected against in transitions in direction *a*, while two genotypes are selected against in transitions in direction *b*.

### 4.2 Model 2: Transitions that change the sex chromosome pair

In this section, we study transitions between male and female heterogamety where, in the course of the transitions, a pair of chromosomes that are initially autosomal are co-opted as new sex chromosomes, while one of the old sex chromosomes becomes autosomal (Figures 2 and 3B).

#### Monte Carlo simulations

Beginning with an XX,AA/XY,AA male-heterogametic system (where X and Y are sex chromosomes and A is an autosome), assume that a mutation appears on an A chromosome, rendering it an A′ such that XX,AA, XX,AA′, XY,AA′, and YY,AA′ individuals are female, while XY,AA and YY,AA individuals are male (Figure 2). If the A′ chromosome reaches sufficiently high frequency in the population, the X chromosome is eliminated, and a YY,AA′/YY,AA female-heterogametic system establishes (direction *a* in Figures 2 and 3B).

We initially assume all six sexual genotypes to be equally fit. Unlike for the case of transitions involving the same chromosome pair, we do not propose a null *expectation* for the fixation probability of the A′ mutation. This is because, if it fixes, it displaces the unlinked X chromosome from the population: this is not the ‘population’ in which the A′ arises, being a mutated A chromosome. (In contrast, in model 1, the X′ chromosome for example is a mutated X chromosome, and if it fixes, it displaces the X chromosome.) Therefore, we shall focus predominantly on comparing the substitution rates of the A′ and X chromosomes (i.e., the substitution rates, respectively, in directions *a* and *b* of Figure 2). We may, however, take as a *reference* neutral fixation probability for both mutations, *ρ*^ref^, that of a mutation of no effect occurring on an autosome: *N ρ*^ref^ = 1/2.

In our simulations, we find the fixation probability of the A′ chromosome to be *Nρ*_A′_ ≈ 1.07 (Figure 6A), substantially higher than our reference value of 1/2. This value is insensitive to the genetic background of the initial mutation (Supplementary Information, Figure S5A). The average conditional fixation time of the A′ in our simulations is approximately 2.65N for each N considered, again regardless of the initial background of the mutation (Supplementary Information, Figure S5B), and again suggestive of drift-like dynamics.

**Figure 6:**
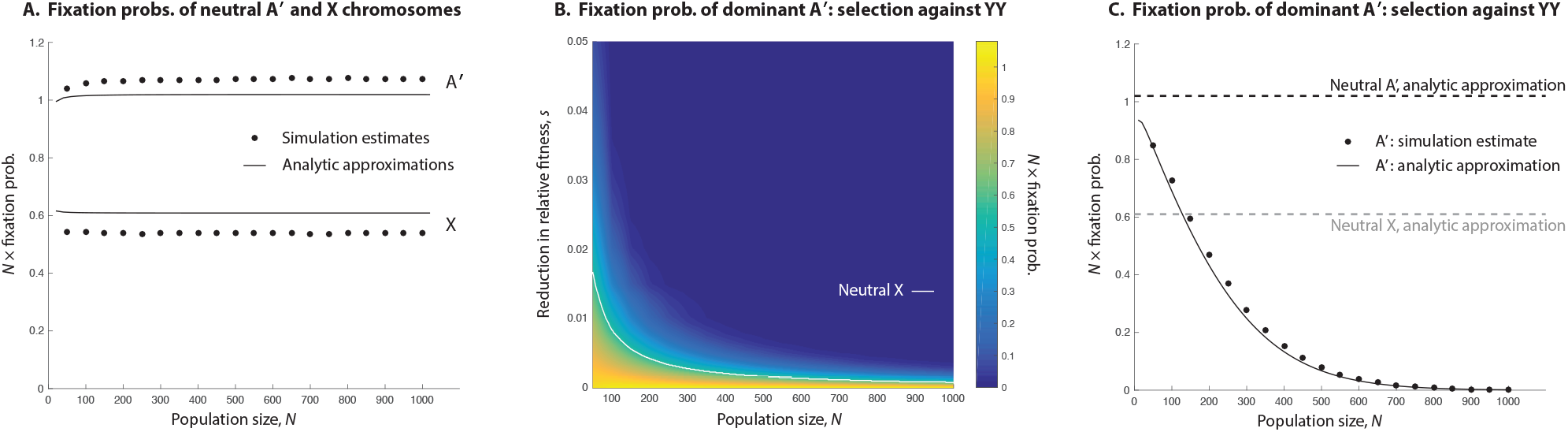
Fixation probabilities and substitution rates in model 2 heterogametic transitions. (A) The fixation probability of a neutral dominant feminizing A′ arising in an XX,AA/XY,AA system is substantially higher than the reverse fixation probability, that of a neutral recessive feminizing X chromosome arising in an YY,AA′/YY,AA system. (B) If YY individuals are selected against (fitness 1 − s) in the transition from XX,AA/XY,AA male heterogamety to YY,AA′/YY,AA female heterogamety (direction *a* in Figure 2), the substitution rate of the A′ chromosome causing this transition is reduced. But selection needs to be sufficiently strong to reduce the substitution rate below that of the neutral X chromosome in the reverse transition (above white line), especially in smaller populations. For ease of visualization, the heat map in panel B is constructed from interpolated data; the raw data are illustrated in Supplementary Information, Figure S7. Panel B is very similar to an analogous heatmap displaying our analytical results (Supplementary Information, Figure S17) (C) Fixation probability of the A′ chromosome when YY individuals have relative fitness reduction *s* = 0.5% (in the Wright-Fisher simulations; the fitness reduction in the Moran analytical results is *s* = 1%, the relevant comparison owing to the two-fold increase in genetic drift under the Moran). Since the reverse transition (direction *b* in Figure 2) does not involve the production of individuals homozygous for the previously sex specific chromosome (the A′), the substitution rates in the two directions are equal where the declining fixation probability of the A′ intersects the flat neutral fixation probability of the X, here at a population size of only about 150 individuals. For larger populations, the recessive X substitutes at a higher rate than the dominant A′.

Turning our attention to transitions in the other direction along the equilibrium path (direction *b* in Figures 2 and 3B), we begin with a female-heterogametic YY,AA′/YY,AA system, and assume that a mutation on a Y chromosome occurs, rendering the chromosome a recessive feminizing X. If this mutation reaches sufficient frequency, the male-heterogametic system XX,AA/XY,AA establishes.

We estimate in our simulations that the fixation probability of this transition is *Nρ*_X_ ≈ 0.54 (Figure 6A), which is fractionally higher than the reference value of 1/2. The background on which the mutation arises has no effect on its fixation probability (Supplementary Information, Figure S6A). The average conditional fixation time of the X chromosome is also close to 2.65N for all values of *N* considered (Supplementary Information, Figure S6B).

To see if there is a directional bias in one direction or the other along the equilibrium path, we assume that these mutations occur at the same rate, *u* per chromosome per generation, and calculate the substitution rates of the two transitions.

The male-heterogametic system XX,AA/XY,AA generates A′ mutations at rate *μ*_A′_ = 2*Nu*, so that the substitution rate from an XX,AA/XY,AA system to a YY,AA′/YY,AA system (in direction *a* of Figures 2, 3B) is about *μ*_A′_*ρ*_A′_ = 2*Nu* × 1.07/*N* = 2.14*u*. Similarly, the female-heterogametic system YY,AA′/YY,AA generates X mutations at rate *μ*_X_ = 2*Nu*, and so the substitution rate from a YY,AA′/YY,AA system to an XX,AA/XY,AA system (direction *b* of Figures 2, 3B) is about *μ*_X_*ρ*_X_ = 2*Nu* × 0.54/*N* = 1.08*u*, or about half that of the reverse transition.

Again, we have assumed no demographic differences between males and females in the above simulations, and so the transitions symmetric to those above (i.e., in directions *c* and *d* in Figure 3B) occur with fixation probabilities and substitution rates equivalent to those we have estimated above (i.e., the fixation probabilities in directions *a* and *b* match those in directions *c* and *d* respectively).

We now consider the role of selective differences between the genotypes. This is an important question here because, unlike in model 1 heterogametic transitions, model 2 transitions are possible without the production of individuals homozygous for a previously sex-specific chromosome. In particular, the transitions in directions *b* and *d* of Figure 3B change the heterogametic system, but do not involve the production of individuals homozygous for the previously sex-specific chromosome. In contrast, the reverse transitions (in directions *a* and *c*) do involve the production of individuals homozygous for previously sex-specific chromosomes. Since we have found these latter transitions, when all genotypes are of equal fitness, to have significantly higher substitution rates than the reverse transitions, we should expect selection to reduce, and when strong enough to overturn, this bias.

We focus on the transition from the male-heterogametic XX,AA/XY,AA system to the female-heterogametic YY,AA′/YY,AA system (direction *a* in Figures 2 and 3B), involving the substitution of a dominant female-determining A′ chromosome as a mutated A. The Y chromosome is sex-specific in the original XX,AA/XY,AA system, and we assume that it has accumulated deleterious recessive mutations such that the two YY genotypes are of fitness 1 − *s*, relative to all other genotypes’ fitness of 1. Figure 6B gives the fixation probability of the A′ chromosome for various values of *s* and *N*. Naturally, the fixation probability decreases as selection against the YY genotypes increases, and this effect is stronger in larger populations [in which selection acts more efficiently (Lanfear et al. 2014)]. Indeed, in large populations, even very small degrees of selection against the YY genotypes are enough to overturn the substitution rate bias in favor of dominant sex determining mutations.

#### Analytical results

We begin by considering the dynamics of the neutral model. Once again, fast-variable elimination can be used to calculate the effective dynamics of the system along the equilibrium path [see Eq. (4)]. For model 2 (as with model 1), 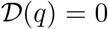 (that is, there is no deterministic contribution to the dynamics along the equilibrium path), but there is a drift-induced selection term, which now takes the form

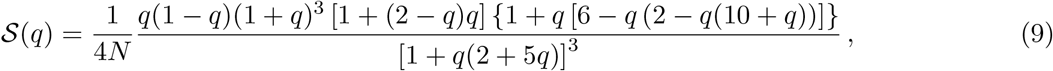

which, along with the expression for diffusion along the equilibrium path,

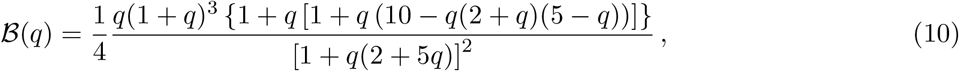

approximates the stochastic dynamics (calculations in Supplementary Information, Section S2.2.2). Recalling that *q* = 0 corresponds to male heterogamety [XX,AA/XY,AA; *p*_1_ = 1/2, *p*_5_ = 1/2 in Eq. (2)], while *q* = 1 corresponds to female heterogamety [YY,AA′/YY,AA; *p*_4_ = 1/2, = 1/2 in Eq. (2)], we find that the drift-induced selection selects for the fixation of female heterogamety *at every point* on the equilibrium path [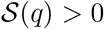 for all *q* ∈ (0, 1)].

As in model 1, we find that demographic fluctuations away from the equilibrium path return to the equilibrium path on average with a bias. It is this bias generates the drift-induced selection term 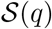. Once again, the drift-induced selection term 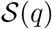 can be split into two components, 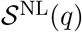 and 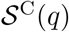, that respectively capture the contribution to 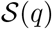 arising from the non-linearity of trajectories taking the system back to the equilibrium path and the curvature of the equilibrium path itself. These terms are plotted in Figure 7A, in which we see that, as with model 1, it is the curvature of the equilibrium path that contributes most to the observed drift-induced selection in model 2.

**Figure 7:**
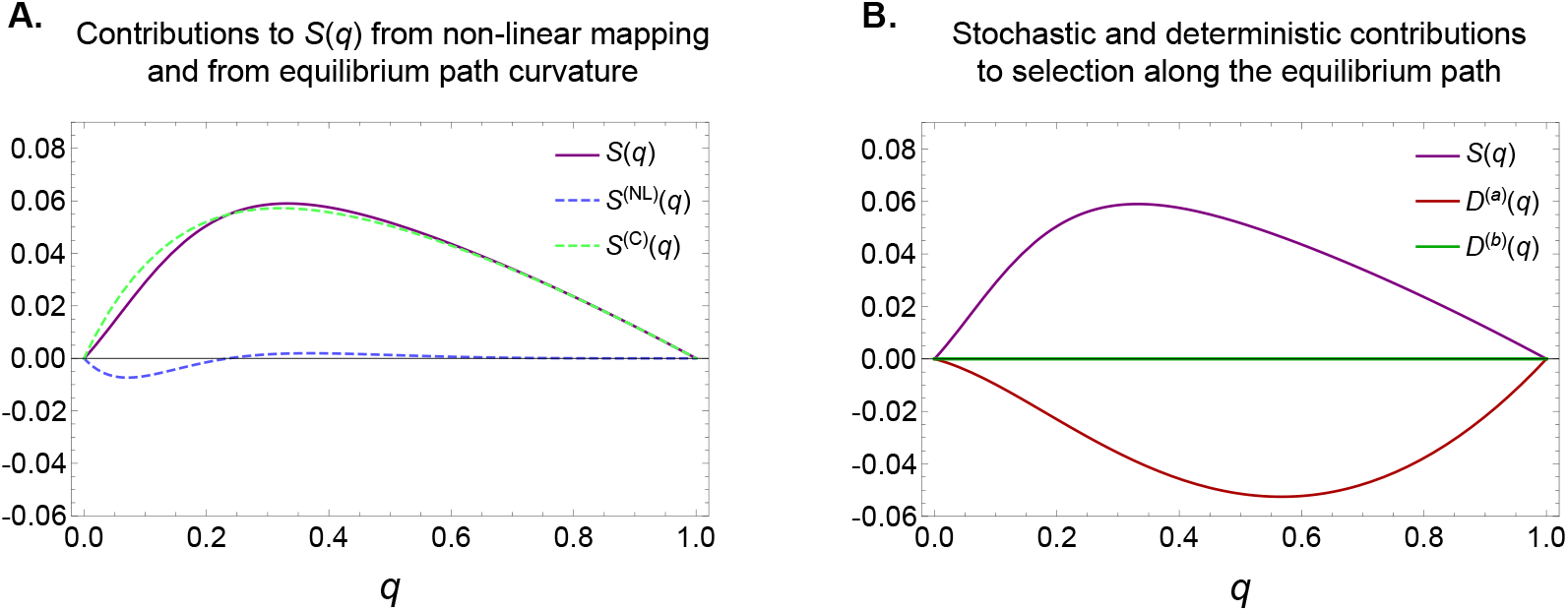
(A) In model 2, as in model 1, random demographic fluctuations induce a selective gradient 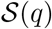 along the equilibrium path in favor of the dominant sex determining chromosome, i.e., causing *q* to increase on average [see Eqs. (4) and (9)]. Again, 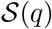 can be divided into two components: 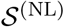, from non-linear components of the mapping of fluctuations back to the equilibrium path, and 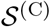, from the curvature of the equilibrium path. As in model 1, curvature of the equilibrium path is the primary contributor to drift-induced selection in favor of increasing *q*. (B) Contributions to average dynamics along the equilibrium path arising from drift-induced selection, 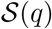, and deterministic selection against individuals homozygous for a previously sex-specific chromosome, 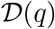 [see Eq. (4)]. Such individuals are only produced in transitions in direction *a* of Figure 2, where the initial population is XX,AA/XY,AA male heterogametic, so that 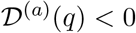 for all *q* ∈ (0, 1) [see Eq. (12)]. In transitions in direction *b* of Figure 2, where the initial population is YY,AA′/YY,AA female heterogametic, no A′A′ individuals are produced, and so 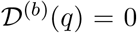 = 0 for all q. In both plots the parameters have been scaled such that *N* = *δ* = 1 in order to facilitate the comparison of contributions to selection along the slow subspace.

Using this single-variable description of the dynamics, the fixation probability for any initial condition *q*_0_ on the equilibrium path can be calculated. In order to determine the fixation probability of mutants in the system, we need to calculate the mapping, for each initial mutation scenario, from *p*_0_ to *q*_0_. For direction *a*, let 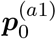 denote the initial conditions when the first A′ chromosome is carried by an XX,AA′ individual, and 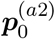 the initial conditions when it is instead carried by an XY,AA′ individual. For direction *b*, let 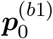 denote the initial conditions when the first X chromosome is carried by an XY,AA′ individual, and 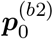 the initial conditions when it is instead carried by an XY,AA individual. Then the respective initial conditions on the equilibrium path can be shown to be (Supplementary Information, Section S2.3.2)

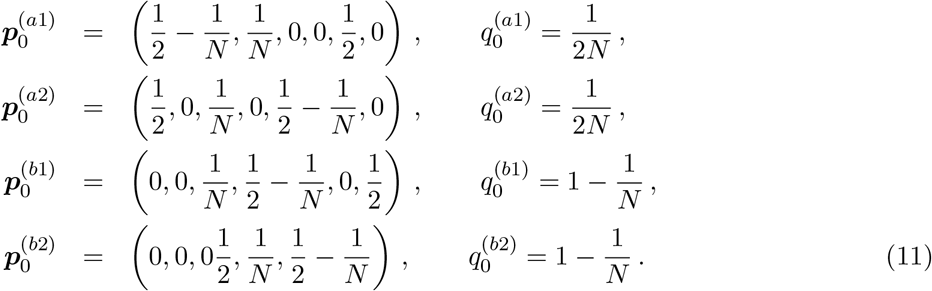

We can now evaluate our numerical expression for the neutral fixation probability at the initial conditions given by Eq. (11). Recall that simulations of the Wright-Fisher process showed the fixation probability of the A′ chromosome to be *Nρ*_A′_ ≈ 1.07, substantially higher than our reference value of 1/2. Our analytical prediction slightly underestimates this fixation probability (*Nρ*_X_ ≈ 1.02; Figure 6A). The background on which the mutation arises has no effect on its fixation probability. This can be understood mathematically by recognizing that the two scenarios initially have an identical component along the slow manifold [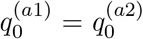 in Eq. (11)]. The computed mean conditional fixation time of the A′ also agrees well with our simulation estimates (Supplementary Information, Figure S5B).

We next consider the fixation probability of an X mutation occurring on an initially autosomal Y. Our Wright-Fisher simulations returned an estimated fixation probability of *Nρ*_A′_ ≈ 0.54. Once again, there is a small discrepancy with our analytical results, which overestimate this value (*Nρ*_X_ ≈ 0.61; Figure 6A). The fixation probability is again the same irrespective of the background on which the X mutation occurs [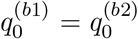 in Eq. (11)]. Again, we may compute the mean conditional fixation time of the X, and find good agreement with our simulation estimates when the variance difference between the Wright-Fisher and Moran processes is accounted for (Supplementary Information, Figure S6B).

We now consider the role of selective differences between the genotypes. As we have noted, comparing transitions in directions *a* and *b*, only those in direction *a* involve the production of individuals homozygous for a previously sex-specific chromosome, and so we focus here on transitions in direction *a*. We assume that the two YY genotypes (with frequencies *p*_4_ and *p*_6_) have elevated death rates Δ_4_ = 1 + *s* and Δ_6_ = 1 + *s* relative to all other genotypes’ death rates of 1. The term 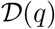 in Eq. (4) now becomes 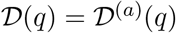, where

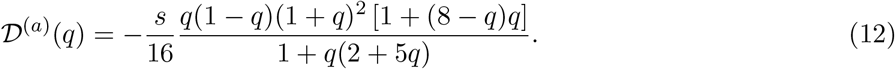

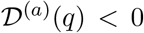 for all *q* ∈ (0, 1), and so this term acts in the opposite direction to 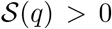 (Figure 7B; calculations in Supplementary Information, Section S2.2.2). There is thus an antagonism between the deterministic contribution to selection 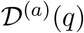 (favoring transitions in direction *b*) and the drift-induced selection (favoring transitions in direction *a*). Which of these dominates the dynamics depends on the strength of selection and the population size; 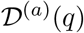 increases with *s*, while 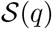 decreases with *N*. Therefore, in large populations, even small degrees of selection against the YY genotypes are enough to overturn the substitution rate bias in favor of dominant sex determining mutations.

In contrast, since transitions in direction *b* do not produce individuals homozygous for a previously sex-specific chromosome, none of the genotypes is selected against. Therefore, 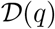 in direction *b* is 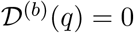.

Whereas in model 1 selection exacerbated the directionality of switching between sex determining systems (while decreasing the overall switching rate), in model 2 we see that deterministic selection and drift-induced selection work in opposite directions. Thus, for small populations with weak selection, transitions in directions *a* and *c* in Figure 3B occur more often than transitions in directions *b* and *d*, while for large populations or strong selection, transitions in *b* and *d* occur more often than transitions in directions *a* and *c*.

## 5 Discussion

We have studied stochastic evolution along two ‘neutral’ equilibrium paths connecting male and female heterogamety. We have shown that, even when all genotypes are equally fit, evolution along these paths is not neutral. Instead, it shows significant substitution rate biases in particular directions, specifically in favor of dominant sex determining mutations. We have demonstrated these biases to be the result of drift-induced selection: random fluctuations off the equilibrium path—which are inevitable in finite populations—return to the equilibrium path with an average directional bias in favor of the dominant segregating sex chromosome. The substitution rates of dominant sex determining mutations that switch the system of heterogamety are, in both of the cases we have studied, substantially higher than those of truly neutral mutations occurring on the same chromosomes.

The system of heterogamety has often been considered an ‘evolutionary trap’ (Bull and Charnov 1985; Rice 1998), owing to the inevitable degradation of the sex-specific chromosome through the operation of Muller’s ratchet (Charlesworth 1978; Charlesworth and Charlesworth 2000). This is known to be at odds with the frequent evolutionary transitions between male and female heterogamety observed and inferred across animals (Hillis and Green 1990; Ezaz et al. 2006, 2009; Pokorna and Kratochvíl 2009; Mank et al. 2006; Mank and Avise 2009; Kaiser and Bachtrog 2010; Bachtrog et al. 2014), and so selective mechanisms have been invoked to explain these transitions [e.g., the operation of sex-specific selection (van Doorn and Kirkpatrick 2010)]. Our demonstration of drift-induced selection in transitions between male and female heterogamety suggests that selective differences between the sex chromosome genotypes may be unnecessary to explain empirical transitions.

On this point, we have shown that small fitness reductions to individuals homozygous for previously sex-specific chromosomes—which are always produced in both model 1 and model 2 transitions involving dominant sex determining mutations—are not enough to overturn the biases caused by drift-induced selection (Figures 4C and 6B). The influence of drift-induced selection in heterogametic transitions is therefore best understood in terms of evolutionary timescale. From an initial heterogametic system with homomorphic sex chromosomes, the sex-specific chromosome gradually accumulates recessive deleterious mutations that would reduce the fitness of individuals homozygous for this chromosome. At some threshold period of time of accumulation of these mutations, the fitness reduction of the homozygote reduces the fixation probability of a dominant sex reversal mutation exactly enough to cancel this mutation’s drift-induced selective advantage (Figures 4C and 6B). Prior to this time threshold, a transition via a dominant sex reversal mutation is likely—it becomes less likely as the threshold is neared. However, progress towards the threshold is reset every time a transition occurs, because each transition creates a new sex-specific chromosome from a chromosome that was previously not sex-specific.

A notable prediction of our findings is that heterogametic transitions should typically involve dominant sex determining mutations, which we have found to enjoy a drift-induced selective advantage over recessive mutations. That is, we should expect transitions usually to occur in directions *a* and *c* in Figures 3A and 3B, and not in directions *b* and *d*. We should note that this general prediction, viz. that the novel sex determining mutations involved in heterogametic transitions should be dominant, is not unique to our theory. For example, under the theory that heterogametic transitions can be driven by linkage between novel sex determining genes and genes with sex-specific fitness advantages (van Doorn and Kirkpatrick 2010), dominance of the sex determining mutation is either required for a transition, or substantially increases the parameter range over which a transition may occur. However our analysis does reveal that this prediction can be recapitulated with a minimal number of biological assumptions.

Evidence in favor of the prediction comes from currently observed intermediate ‘multi-factorial’ systems (Bachtrog et al. 2011). Consider the well-known example of the platyfish *Xiphophorus maculatus*, which has a system of sex determination that is intermediate between male and female heterogamety: females are XX, WX, or WY, while males are XY or YY (Kallman 1965, 1968). In principle, such a system could have arisen either in directions *a* or *b* of Figure 3A, depending on the ancestral heterogametic system. Mapping known systems of heterogamety in the genus *Xiphophorus* (Tree of Sex Consortium 2014) onto a phylogeny of the clade (Cui et al. 2013) suggests this ancestral system to be male heterogamety, in which case the intermediate system of *X. maculatus* has arisen in direction *a* of Figure 3A, via a dominant sex reversal mutation, consistent with our prediction. The western clawed frog, *Xenopus tropicalis*, also has an intermediate system: females are ZW or WW, and males are ZZ, ZY, or WY (Roco et al. 2015). This system could have arisen in direction *c* or *d* in Figure 3A, depending on whether female or male heterogamety is ancestral, respectively. The mechanism of sex determination has yet to be determined for most members of the genus *Xenopus* (Tree of Sex Consortium 2014; Roco et al. 2015), but the well-studied *X. laevis* is female-heterogametic (Chang and Witschi 1956), as is *X. borealis* (Furman and Evans 2016). If female heterogamety is ancestral to the intermediate system of *X. tropicalis*, then this intermediate system would have arisen in direction *c* of Figure 3A, again consistent with our prediction. Though it is possible that balancing selection operates to stabilize these observed instances of multi-factorial systems [e.g., Orzack et al. (1980)]—with the important suggestion that, because observed instances are rare, most intermediate multi-factorial systems are transitional—the drift-induced selection that we have discovered operating at all points on the slow subspace near the line of equilibria will, even in this case, act so as to make invasion of dominant sex determining mutations more likely.

The prediction that transitions should usually involve dominant sex determining mutations can also be tested by reciprocal crosses of species or populations on either side of a recent heterogametic transition, if the ancestral system is known. In the frog *Rana rugosa*, populations in northern Japan are female-heterogametic, while those in southern Japan are male-heterogametic (Nishioka et al. 1993; Miura et al. 1998). The sex chromosomes in these populations are all homologous (Uno et al. 2008), and so a model 1 transition appears to have occurred. Because the ancestral system is male heterogamety (Ogata et al. 2003), the candidate directions in Figure 3A are *a* and *d*. These two directions can be distinguished by crossing a homogametic male (from the north) with a homogametic female (from the south). If the transition occurred in direction *a*, all the offspring from this cross should be male, but if it occurred instead in direction *d*, all the offspring should be female. This test has been carried out using homogametic males from Hirosaki (in the north) and homogametic females from Kumano (in the south), the reciprocal cross of which yielded almost all male offspring (Nishioka et al. 1993). Again, this is consistent with our prediction. [Crossing heterogametic males and females yielded a sex ratio of 1/2 (Nishioka et al. 1993), consistent with the model 1 transitions we have studied, though not informative of which direction the transition was in.]

We have referred throughout to ‘sex chromosomes’, but in reality we are talking about genes of major sex determining effect, the presence or absence of which acts as a switch to direct development down separate molecular pathways, or sex determining ‘cascades’, which then produce males and females. The downstream components of these cascades tend to be widely conserved (Beukeboom and Perrin 2014), but there is significant lability in the upstream components—through the addition of new sex determining genes to the top of the cascade (Wilkins 1995) and the shuffling of genes already in a cascade (Schartl 2004)—suggesting that these cascades evolve ‘from the bottom up’ (discussed below). In particular, the gene at the very top of the cascade, i.e., the major sex determining gene, varies widely across and within clades (Beukeboom and Perrin 2014). These observations of sex determining cascades are consistent with the predictions of our model. Our model predicts successive transitions involving dominant sex determining mutations, with comparatively few reversals involving fixation of recessive sex determining mutations, and so we expect either the expansion of sex determining cascades, or their shuffling, but seldom their contraction.

We have found that dominant sex determining mutations fix with higher probability than comparable truly neutral mutations, and with higher probability than the reverse recessive mutations, but also that their fixation probabilities are proportional to 1/*N*, and therefore decrease as population size gets large. It might be objected that their neutral fixation is therefore not a relevant possibility in large populations. As with the study of neutral substitutions elsewhere in the genome, this depends on how often the relevant mutations are produced: if the mutation rate to new dominant sex determining genes (including changes in the locus of the major sex determining gene) is sufficiently high, then their fixation, even if its probability scales with 1/*N*, is empirically relevant. There are two reasons to believe this condition to hold in the case of sex determining mutations: In principle, the many ways in which such mutations can arise, and in practice, the heterogeneity in major sex determining genes observed between and within clades.

The generation of new sex determining genes or loci is known to occur through many possible paths, in addition to standard sequence-mutation events: (i) translocation of a gene in a sex determining pathway and a resulting shift in expression or function (Traut and Willhoeft 1990; Charlesworth et al. 2005), (ii) duplication of such a gene and the subsequent acquisition of new function (Schartl 2004; Bewick et al. 2011) (iii) mutation of a a major sex determining factor’s regulatory elements, such as transcription factors (Takehana et al. 2014; Beukeboom and Perrin 2014). All of these possibilities are more likely to result in the creation of a new major sex determining gene if they occur in a gene already involved in the sex determination pathway. These additional sources of mutation exacerbate the shuffling of gene cascades and the general movement of genes up the cascade (Schartl 2004).

An excellent example of this phenomenon is seen the DMRT1 gene, of the DM domain family of transcription factors (Matson and Zarkower 2012; Beukeboom and Perrin 2014). DMRT1 is currently thought to be the major sex-determining factor in birds, but is also an upstream component of the SOX-9 feedback loop early in the mammalian sex determination cascade (the major sex determining gene of which is SRY) (Beukeboom and Perrin 2014). DMRT1 is involved in downstream testicular differentiation in amphibians, but a duplicated homolog, DM-W, is the major sex determining factor in some species of *Xenopus* (Shibata et al. 2002; Yoshimoto et al. 2008; Bewick et al. 2011). In fact, seven paralogs of DMRT1 are known in mammalian lineages alone, and many more are common across vertebrates, although their function in sex determination and differentiation diverges widely (Herpin and Schartl 2011; Matson and Zarkower 2012; Beukeboom and Perrin 2014). This conserved element of the sex-determination cascade in vertebrates illustrates not only the shuffling of genes in these cascades, but also the recurrent evolution of new sex determining mechanisms through structural mutation.

In comparing the substitution rates of dominant and recessive sex determining mutations, we have assumed that their respective mutation rates, i.e., the rates at which they are generated, are equal. It is possible that this is not the case, and that one class of sex determining mutations are generated more rapidly than the other (Hillis and Green 1990; Bachtrog et al. 2011). If this were the case, it would simply be a distinct mechanism by which we expect one class of sex determining gene to be more prevalent than the other. We should note that this is not as simple as comparing the rates of generation of gain-of-function and loss-of-function mutations. Suppose, for example, that a system is initially XX/XY male heterogametic, and consider a transition in direction *a* of Figure 1A, involving fixation of a dominant feminizing X′ mutation. Depending on the molecular functioning of the initial XX/XY system, this dominant X′ could be gain-of-function or loss-of-function. If the Y is initially dominant male-determining [as, e.g., in mammals (Koopman et al. 1991; Beukeboom and Perrin 2014)], then a mutation on the X chromosome that blocks the Y’s activity (a gain-of-function mutation) would be dominant feminizing, as we require. If, however, the initial system depends on the ratio of some gene (or genes) on the X chromosome with respect to autosomes [as in *Drosophila* (Bridges 1921; Beukeboom and Perrin 2014)], then a loss-of-function mutation to a relevant gene on the X chromosome could be dominant feminizing.

We have identified the force driving our results to be drift-induced selection operating along the equilibrium path (in the neutral case) or in its near vicinity (in the case with selection). While drift-induced selection as a force in natural selection has analogs that have been known for some time [e.g., Gillespie’s criterion (Gillespie 1974, 1977)], the demonstration that it acts endogenously in systems of biological interest is relatively recent (Parsons et al. 2010; Lin et al. 2012; Kogan et al. 2014; Constable et al. 2016; Newberry et al. 2016; Chotibut and Nelson 2017). We suspect that drift-induced selection will come to be recognized as an important force in many dynamical systems in population biology.

## 6 Acknowledgements

We are grateful to David Haig, Dan Hartl, Sally Otto, and Naomi Pierce for helpful comments. The computations in this paper were run on the Odyssey cluster supported by the FAS Division of Science, Research Computing Group at Harvard University. PM is supported by an NSF graduate research fellowship. GWAC thanks the Finnish Center for Excellence in Biological Interactions and the Leverhulme Early Career Fellowship provided by the Leverhulme Trust for funding.

